# Recombinant Expression of Aldehyde Dehydrogenase 2 (ALDH2) in *Escherichia coli* Nissle 1917 for Oral Delivery in ALDH2-Deficient Individuals

**DOI:** 10.1101/674606

**Authors:** Tim Ho, Catherine Chang, Justin Wu, Iris Huang, Leona Tsai, Justin Lin, Emily Tai, Caroline Chou, Justin Yang, Yvonne Wei, Catherine Yeh, William Chen, Dylan Lu, Charlotte Chou, Longan Su, Nicole Chang, Colin Huang, Chloe Wang, Paul Abrena, Christy Cheung, Cassandra Yeh, Shantih Whiteford, Phoebe Chen, Austin Huang, Aire Wu, Benjamin Wei, Eugene Kao, Nicholas Lin, Anna Chang, Jake Yang, Yasmin Lin, Sean Tsao, Nicholas Ward, Teresa Chiang, Jude Clapper

## Abstract

Turning red after consuming alcohol may seem like a mere social inconvenience. Yet, this flushing response is caused by an accumulation of acetaldehyde, a carcinogenic intermediate of alcohol metabolism. Aldehyde dehydrogenase 2 (ALDH2) deficiency, the result of a point mutation, produces a less efficient ALDH2. The resulting accumulation of acetaldehyde greatly increases the risk of developing esophageal and head and neck cancers. In this study, we produced recombinant ALDH2 in the probiotic *E. coli* Nissle 1917, which successfully reduces acetaldehyde levels in simulated oral conditions. Packaged in a hard candy, the ALDH2-probiotic would remain in the mouth to specifically target salivary acetaldehyde. Using mathematical modeling, we also determined how much recombinant ALDH2 is needed to reduce elevated acetaldehyde levels.

**Financial Disclosure:** This work was funded by Taipei American School. The funders had no role in study design, data collection and analysis, decision to publish, or preparation of the manuscript.

**Competing Interests:** The authors have declared that no competing interests exist.

**Ethics Statement:** N/A

**Data Availability:** Yes – all data are fully available without restriction. Sequences for the plasmids used in this study are available through the Registry of Standard Biological Parts. Links to raw data are included in Supplementary Information.

## Introduction

### ALDH2 Deficiency

ALDH2 deficiency, more commonly known as Alcohol Flushing Syndrome or Asian Glow, is a genetic condition that interferes with the metabolism of alcohol. Normally, ethanol is first converted to acetaldehyde (a toxic intermediate) by the enzyme alcohol dehydrogenase (ADH). Then, acetaldehyde is converted to acetate, which can be safely metabolized in the body by aldehyde dehydrogenase 2 (ALDH2) (Figure 1). People with ALDH2 deficiency, however, have a point mutation which leads to a less efficient mutant (ALDH2*2), with lower binding affinity to the coenzyme NAD+ [1], [2]. Enzymatic activity in ALDH2-deficient individuals can be as low as 4% compared to wild type ALDH2*1 activity [2], [3], [4], [5]. As a result, acetaldehyde accumulates and induces an inflammatory response that causes the skin to flush or appear red after drinking alcohol [6]. Facial flushing is the most obvious result of ALDH2 deficiency, but symptoms also include headaches, dizziness, hypotension, and heart palpitations [3], [7].

**Figure 1.**
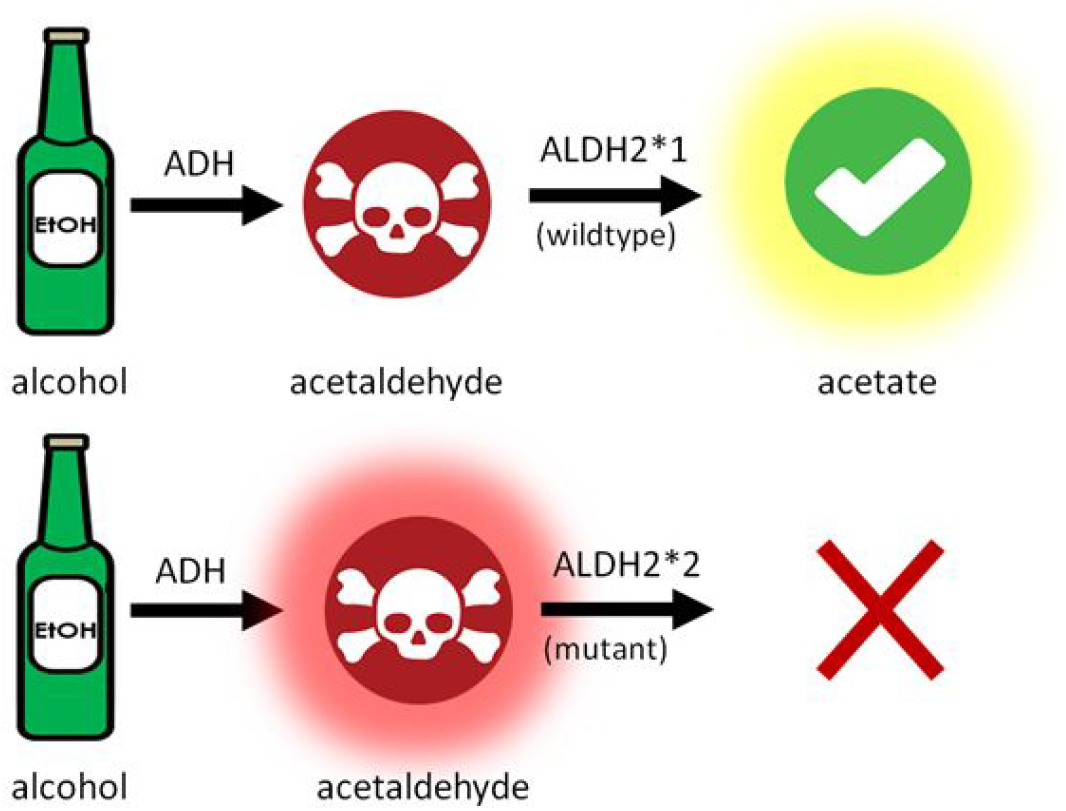
Acetaldehyde accumulates in ALDH2-deficient individuals. Ethanol is first converted to a toxic intermediate, acetaldehyde, by alcohol dehydrogenase (ADH), then converted to acetate by wild type ALDH2*1. The mutant form, ALDH2*2, cannot fully convert acetaldehyde into acetate, and acetaldehyde accumulates as a result.

ALDH2 deficiency affects 540 million people—8% of the world population. In East Asia (which includes Japan, China, and Korea), this is a much bigger problem, where the number rises to 36% [8]. In our home, Taiwan, approximately 47% of the population carries this genetic mutation—the highest ALDH2-deficient percentage in the world [9].

### Health Concerns & Risks

Turning red after consuming alcohol may seem like a mere social inconvenience. Yet, behind this red complexion lies a far more serious problem. The **International Agency for Research on Cancer classifies acetaldehyde associated with alcohol consumption as a Group 1 carcinogen** [10]. Alcohol and acetaldehyde have been shown to reduce thymidine incorporation into DNA, interfering with DNA synthesis [11], [12]. In other studies with human lymphocyte cells, acetaldehyde increased chromosome damage or the frequency of sister-chromatid exchanges [13], [14], [15]. When rats were exposed to acetaldehyde, inhalation led to abnormal changes in the nasal mucous membrane, such as degeneration of the tissue, increased cell proliferation, and the development of carcinomas [16].

Acetaldehyde levels over 50 μM are considered toxic and cause mutations in DNA [17], [18]. In human studies where roughly 2 to 3 servings of alcohol (0.5-0.6 g alcohol/kg body weight) were ingested, salivary acetaldehyde levels in some individuals reached over 100 μM, compared to normal levels of <20 μM without drinking [19], [20], [21].

For people who carry normal, functional ALDH2*1, acetaldehyde can quickly get broken down (Figure 2). For those who are ALDH2-deficient *and* drink, however, acetaldehyde can accumulate to toxic levels. The strongest effects are seen in the mouth. Studies show that after alcohol consumption, **salivary acetaldehyde levels are significantly higher than blood acetaldehyde levels** [20], [22]. Since there are other aldehyde dehydrogenases, such as ALDH1A1, throughout the body to help metabolize acetaldehyde, blood acetaldehyde concentrations remain relatively low [23]. ALDH2 levels are also much higher in the liver, which can break down most acetaldehyde before it enters the bloodstream [24]. This may explain why ALDH2 deficiency does not seem to be directly associated with the development of liver, breast, colorectal, or other common alcohol-related cancers [25], [9]. Instead, the effects of ALDH2 deficiency are mainly localized in the upper digestive and head and neck region.

**Figure 2.**
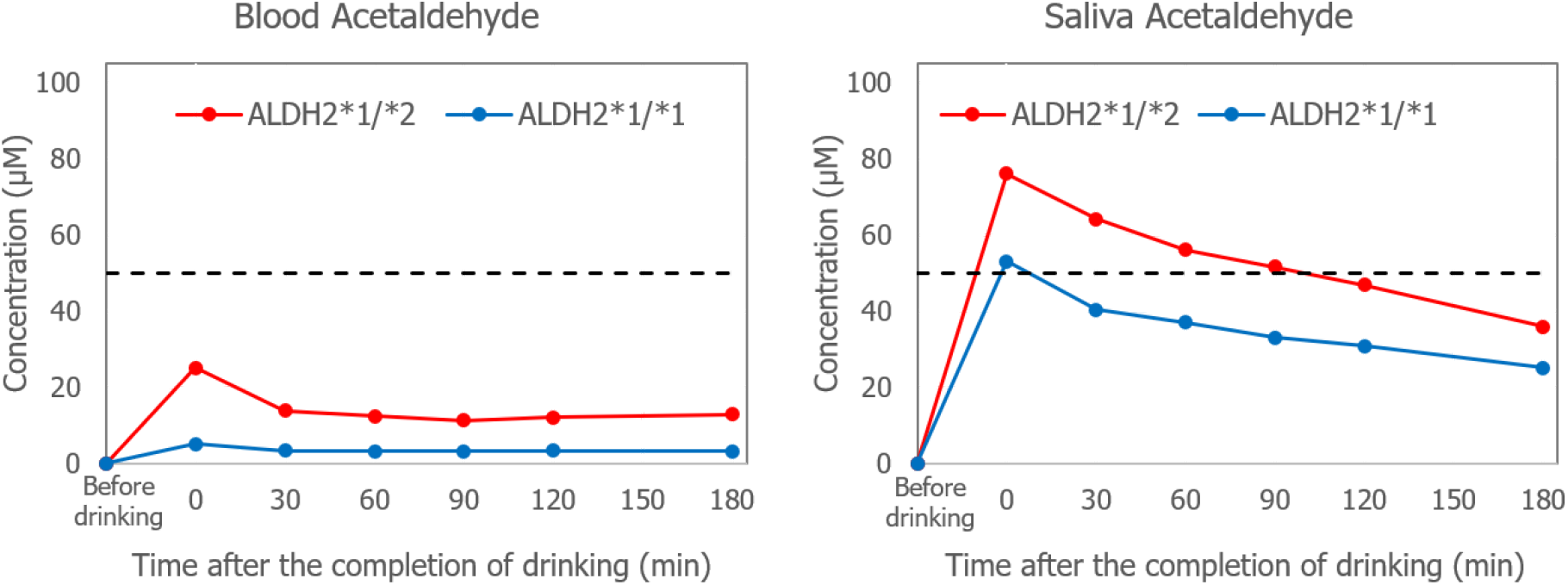
Salivary acetaldehyde levels are much higher in heterozygous (ALDH2*1/*2) individuals, compared to homozygous wild type (ALDH2*1/*1) individuals. Salivary and blood acetaldehyde levels were recorded for individuals before and after drinking 0.6 g ethanol/kg body weight. Salivary acetaldehyde levels are also significantly higher than blood acetaldehyde after drinking (figure adapted from Yokoyama *et al.*, 2008) [20]. Acetaldehyde levels over 50 μM are considered toxic (dashed line).

### Heightened salivary acetaldehyde levels increase the risks of developing esophageal and head and neck cancers

*In vivo* studies with rats have shown that acetaldehyde administered orally has a tumor-promoting effect (abnormally fast cell growth and division) on the upper respiratory-digestive tract [26]. Depending on how much alcohol is consumed, people with ALDH2 deficiency are two to eight times more likely to develop head and neck cancers (including oral cancer, pharyngeal cancer, laryngeal cancer, etc.), and two to twelve times more likely to develop esophageal cancer compared to people with normal ALDH2*1 [27], [28], [29], [30], [31], [32], [33], [34], [35].

Today, it is common to simply use cosmetics to conceal the flushing, or take antihistamines to prevent the release of histamines and the resulting redness [36], [37], [38], [39]. These solutions, however, do not actually address the buildup of acetaldehyde, the main contributor to increased esophageal and head and neck cancers risks [40], [41]. On the contrary, these options may actually increase health risks as they conceal the visible symptoms without reducing acetaldehyde levels [8]. Other treatments include antioxidants that may directly interact with acetaldehyde, but these mainly reduce acetaldehyde accumulation in the blood, instead of treating the acetaldehyde accumulation in saliva [41], [42].

To directly address the increased esophageal and head and neck cancer risks, we aim to reduce salivary acetaldehyde levels. Here, we describe testing and delivery of recombinant ALDH2*1 as a candy to maintain normal acetaldehyde levels in the mouths of ALDH2-deficient individuals.

## Materials and Methods

### Construct Design

#### ALDH2*1 and ALDH2*2

Human ALDH2*1 (iGEM Parts Registry: BBa_M36520; NM_001204889.1) and human ALDH2*2 was modified to remove an internal PstI site and lower the guanine and cytosine content (BBa_K2539150), as recommended by Integrated DNA Technology (IDT). Mutant ALDH2*2 consists of a single G to A point mutation (BBa_K2539250) which leads to a E487K change in the protein. ALDH2 sequences were flanked by an upstream strong promoter and strong ribosome binding site (RBS) combination (BBa_K880005), and a downstream transcription terminator (BBa_B0015) to maximize expression. DNA was synthesized by IDT (BBa_K2539100 and BBa_K2539200), cloned into the standard pSB1C3 Biobrick backbone, and sequenced by Tri-I Biotech.

#### His-tagged ALDH2*1 and ALDH2*2

For protein purification, an additional N-terminal HIS-tag (6xHIS) was added to the start codon of both ALDH2*1 and ALDH2*2 sequences. DNA was synthesized by IDT (BBa_K2539101 and BBa_K2539201), cloned into the standard pSB1C3 Biobrick backbone, and sequenced by Tri-I Biotech.

### Protein Purification

To purify the HIS-tagged proteins, 150 mL of bacterial culture was centrifuged, lysed, and filtered before the cell extract sample was run through a nickel column (GE Healthcare, 11-0033-99). A low-concentration imidazole buffer was run through the column to wash out any non-specific proteins. Finally, we eluted and collected the HIS-tagged ALDH2 proteins by running a high-concentration imidazole buffer through the column (imidazole competes with the HIS tag to bind with nickel ions in the column) [43]. SDS-PAGE was used to check protein content at different steps of the purification process: lysed cell sample, flow-through after the wash buffer, and final eluate containing the purified protein.

### Artificial Saliva

To simulate oral conditions, enzymatic activity tests were run using artificial saliva. We chose to prepare artificial saliva using Biotene Dry Mouth OralBalance Gel, a concentrated gel used to treat xerostomia (more commonly known as dry mouth). The gel contains many of the same ingredients as artificial saliva, such as cellulose, glycerin, and xylitol [44] [45] [46]. The mean viscosity of saliva at room temperature is 2.1 centipoise (cP), and the mean viscosity of water at room temperature is 0.95 cP [47], [48]. We diluted the gel so that the resulting solution would be about 2 times as viscous as water, and then calculated the viscosity of our solutions by adapting the ball drop procedure from Tang (2016) [48]. Diluting 6.5 grams of Biotene gel in 50 mL of water produces a solution with a viscosity of 2.09 cP, which closely mimics saliva.

### ALDH2 Enzyme Activity Assay

The reduction of NAD+ to NADH occurs when ALDH2 converts acetaldehyde into acetate. This reaction takes place in a 1:1 ratio, so production of NADH can be used to test ALDH2’s ability to metabolize acetaldehyde. We used reagents from a kit (Megazyme, K-ACHYD) and quantified the amount of NADH produced by taking absorbance readings at 340 nm. This wavelength is highly absorbed by the reduced form, NADH, but not the oxidized form, NAD+ [49], [50].

We calculated acetaldehyde concentration using the following equation:

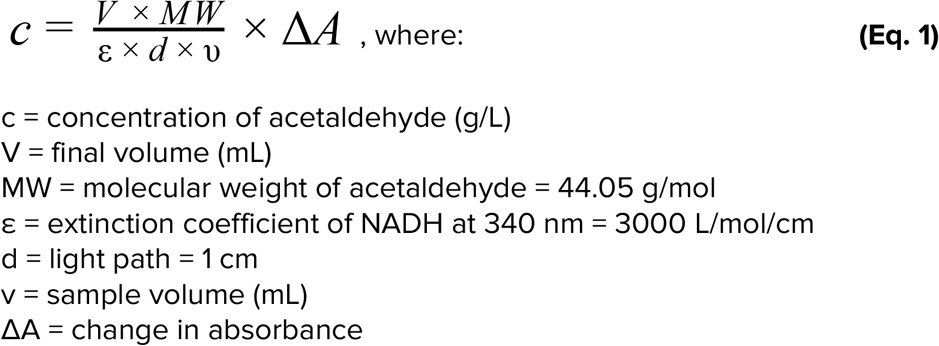

#### Testing Cell Lysate

3 mL bacterial cultures were prepared, and the populations were standardized: the absorbance at OD600 of each culture was measured and cultures were diluted accordingly to achieve the same population in all samples. The cultures were spun down, and the pellets were resuspended and lysed using xTractor Lysis Buffer (Takara Bio, 635671). After lysis, 500 µL of supernatant was transferred into quartz cuvettes containing a pH 9.0 buffer, distilled water, and excess NAD+ (solutions were provided by the Megazyme kit). 50 µL of 0.4 M acetaldehyde was added to initiate the reaction, and absorbance values at 340 nm were recorded for 40 minutes. For all functional tests, the acetaldehyde to water ratio was 1:59 (the total volume was 2.95 mL).

#### Testing Purified HIS-tagged ALDH2

The procedure described above was followed, except that instead of cell lysates, 5 µg of purified ALDH2 was added into quartz cuvettes containing a pH 9.0 buffer, distilled water or artificial saliva, and excess NAD+.

### *E. coli* Nissle 1917 (*Ec*N) Transformation

We obtained a sample of *Ec*N (Mutaflor) from Pharma-Zentrale GmbH. Bacterial cultures were prepared using LB broth, and the cells were made competent by treatment with calcium chloride. GFP- and ALDH2-expressing plasmids were then chemically transformed into competent *Ec*N following standard *E. coli* protocol.

### ALDH2 Probiotic Candy Prototype

10 mL of probiotic *Ec*N culture was pelleted and resuspended in 10 mL distilled water. To determine the heat threshold of *Ec*N, the probiotic mixture was heated from 25°C to 80°C, and 3 µL of bacteria was plated at 5°C intervals. To make our own engineered *Ec*N candy, 30 g of rock sugar and 30 g of gelatin were added to 120 mL of water (Figure 3). The mixture was heated to about 70°C, or until all of the ingredients fully dissolved. GFP-, ALDH2*1-, and ALDH2*2-*Ec*N were grown, pelleted, resuspended in 50 µL of water, and added to the gelatin/candy mixture after it cooled down to 55ºC. 3 mL portions were pipetted into ice cube trays and left to solidify and dry over a period of three days.

**Figure 3.**
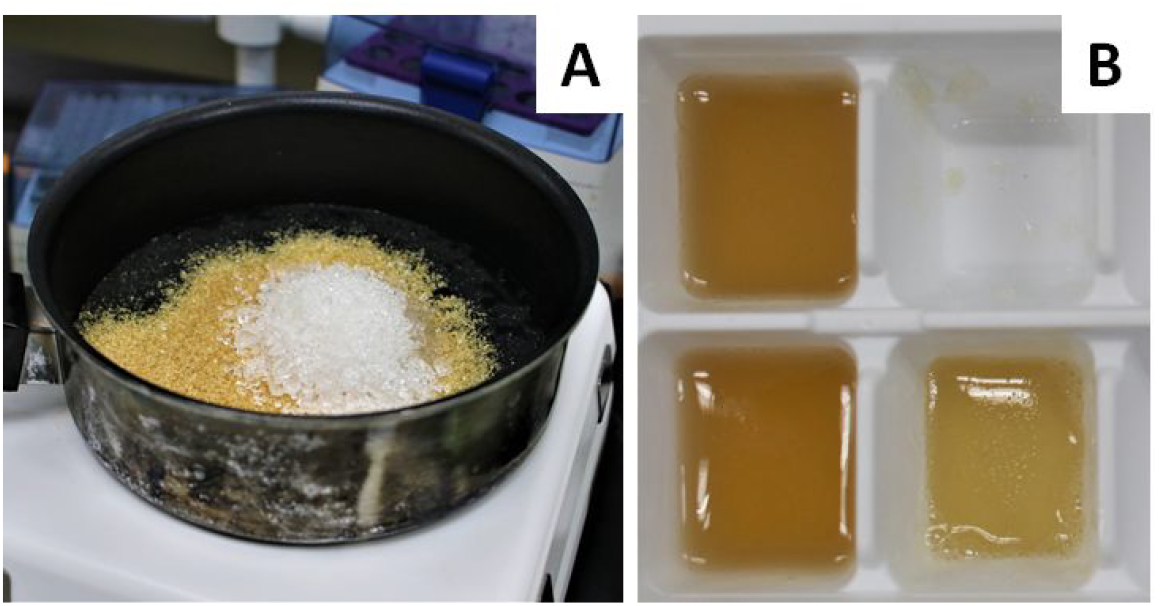
Making *Ec*N probiotic candy. **(A)** 30 g of rock sugar and 30 g of gelatin were dissolved in 120 mL of water. **(B)** After heating, 3 mL of the probiotic candy mixture was pipetted into each ice cube tray and let cool.

#### Testing EcN Candy

We compared the dissolution rate of our *Ec*N candy with commercial lozenges (Ricola and Nin Jiom herbal candies) by gently stirring them in artificial saliva at 37°C. We also determined that commercial lozenges dissolve in the mouth after an average of 13 minutes. To test the candy’s ability to metabolize acetaldehyde, it was dissolved in artificial saliva at 37°C, centrifuged to isolate ALDH2-*Ec*N, and then lysed. Cell lysates were tested using artificial saliva at 37°C.

## Results & Discussion

### Purified ALDH2*1 Metabolizes Acetaldehyde Faster than ALDH2*2

#### Construct Design, Expression, and Protein Purification

HIS-tagged ALDH2*1 and ALDH2*2 expression constructs were assembled (BBa_2539101 and BBa_2539201, respectively, and confirmed by PCR check as well as sequencing (Figure 4). SDS-PAGE was used to check protein content at different steps of the purification process: lysed cell sample, flow-through after the wash buffer, and final eluate containing the purified protein (Figure 4). HIS-tagged ALDH2*1 should be around 56 kDa. In the initial cell lysate lane, there was a band around 56 kDa (Figure 4). This band disappeared in the wash buffer flow-through lane, and reappeared in the eluate. A similar band around 50 kDa was also seen in the eluate when we purified ALDH2*2 (Figure 4). For both HIS-tagged samples, there were other proteins present in the eluate, but the gels showed that we now have purer samples of both HIS-tagged ALDH2*1 and ALDH2*2.

**Figure 4.**
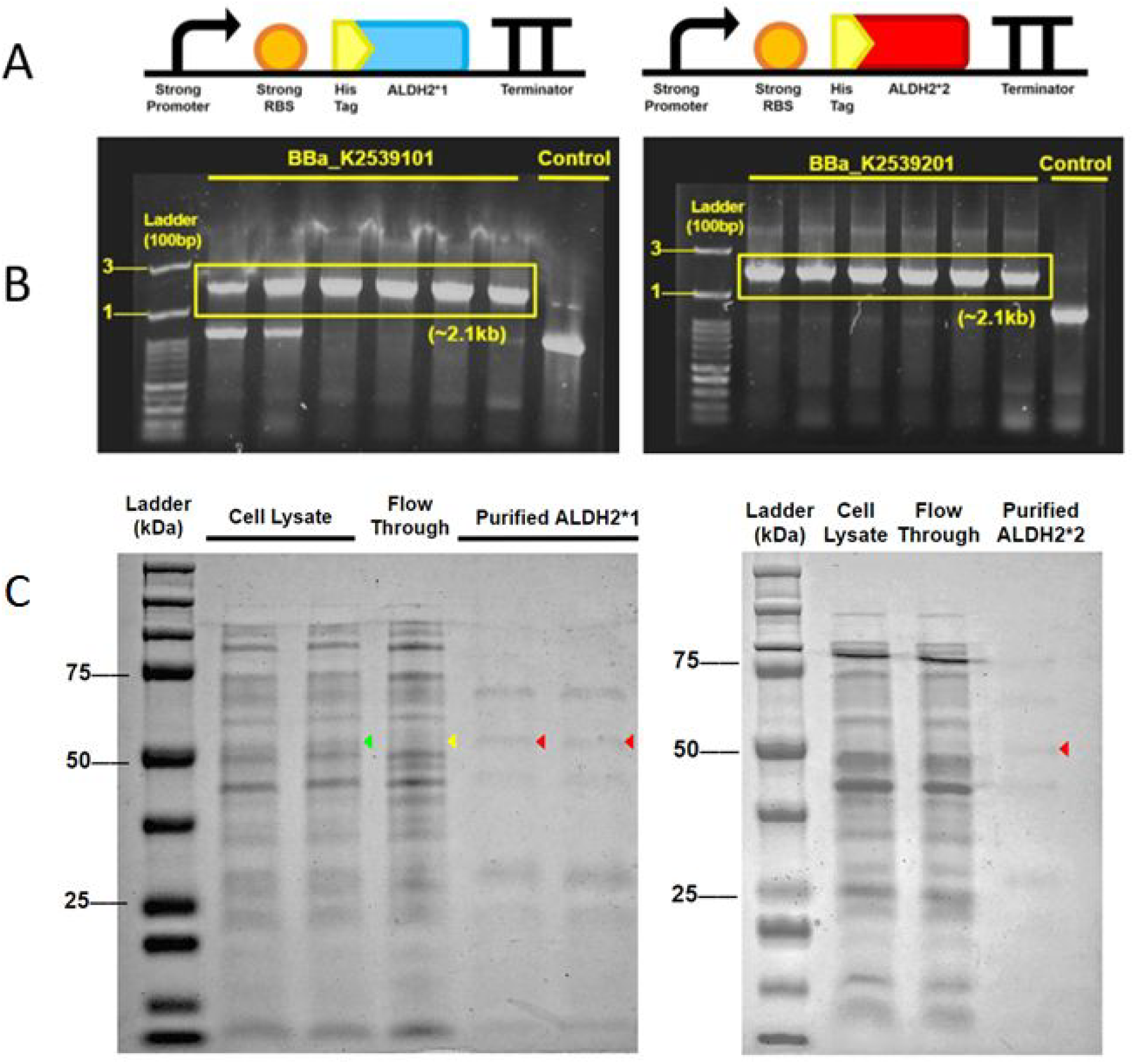
Design, expression, and purification of HIS-tagged ALDH2*1 and ALDH2*2. **(A)** DNA constructs (BBa_K2539101 and BBa_K2539201) contain a constitutive strong promoter, strong RBS, a start codon with a 6x HIS tag, human ALDH2*1 or ALDH2*2 (respectively), and a transcription terminator. **(B)** PCR check for both constructs using VF2 and VR primers produced at the expected size of 2.1 kb (yellow box). **(C)** SDS-PAGE results show protein content at different steps of protein purification. **Left**: A band around 56 kDa in the cell extract (green) and the eluate (red), but not present in the wash buffer flow through lane (yellow), matches our expected HIS-tagged ALDH2*1. **Right:** The same band can be seen in the eluate lane for HIS-tagged ALDH2*2 (red).

#### Measuring ALDH2 Enzyme Activity

To test the enzymatic activity of our recombinant ALDH2, we measured the amount of NADH produced when acetaldehyde is converted to acetate (Figure 5). NADH was quantified by taking absorbance readings at 340 nm. This wavelength is highly absorbed by the reduced form, NADH, but not the oxidized form, NAD+ [49], [50]. Both wild type ALDH2*1 and mutant ALDH2*2 should metabolize acetaldehyde and produce NADH as a result, but ALDH2*1 should be more efficient compared to ALDH2*2. To simulate oral conditions, all tests were run with artificial saliva and at body temperature. Our results show that while both forms of the enzyme can metabolize acetaldehyde, **ALDH2*1 produced significantly higher NADH levels (by about 3.5 times) compared to ALDH2*2** (Figure 5).

**Figure 5.**
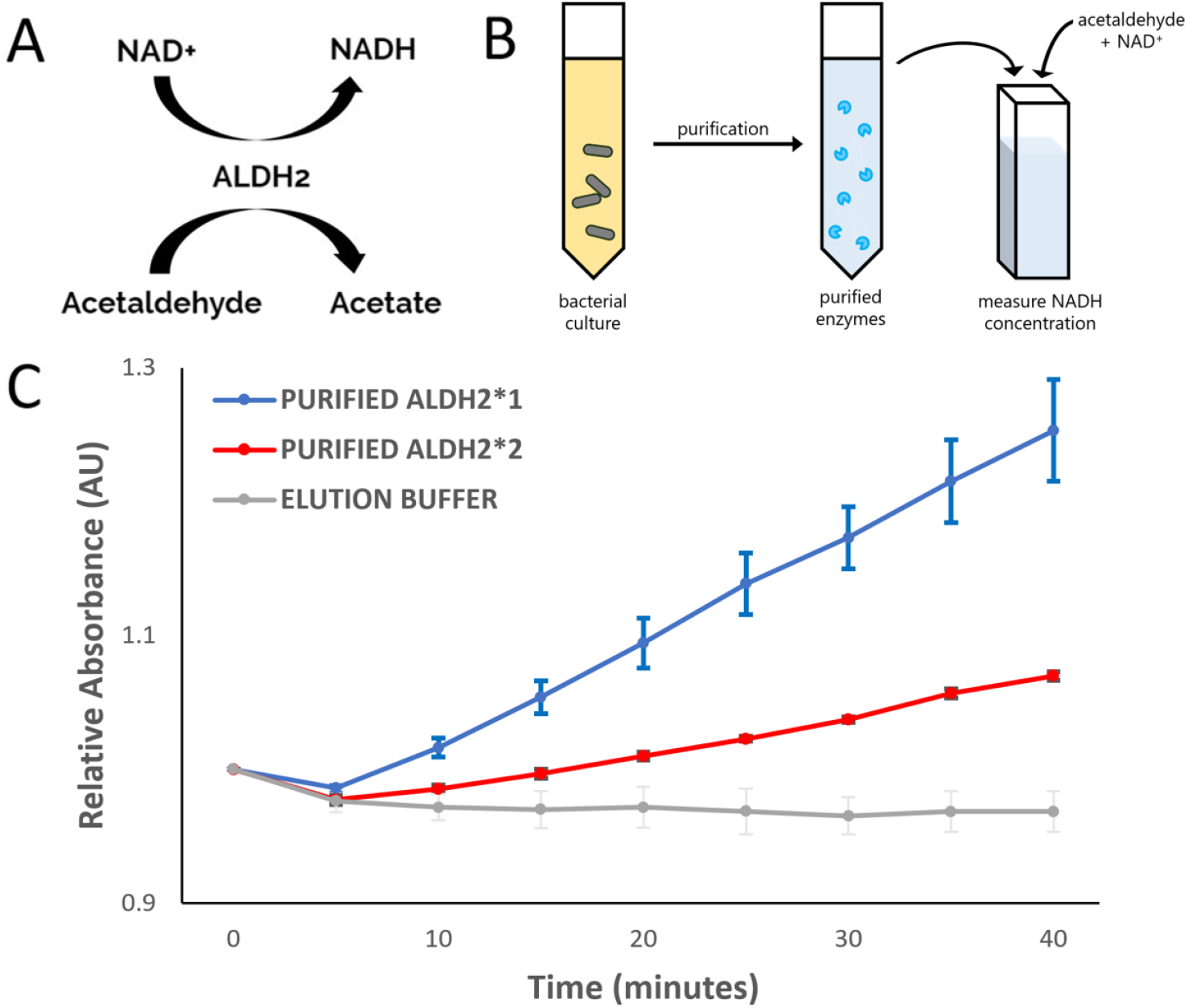
Wild type ALDH2*1 metabolizes acetaldehyde significantly faster compared to mutant ALDH2*2 under simulated oral conditions. **(A)** The conversion of acetaldehyde to acetate by ALDH2 uses NAD+ and produces NADH. **(B)** Experimental setup. Purified enzyme samples were mixed with acetaldehyde and NAD+ to initiate the reaction. NADH concentration was measured by taking absorbance readings at 340 nm. **(C)** Enzymatic activity of purified ALDH2*1 and ALDH2*2 was tested in artificial saliva at 37°C. A negative control containing only elution buffer (from the protein purification process) was also included (gray). Under these conditions, ALDH2*1 metabolized significantly more acetaldehyde compared to both ALDH2*2 and the negative control. Error bars represent standard error.

### ALDH2 Delivery Using the Probiotic *E. coli* Nissle 1917 (*Ec*N)

In a survey (n = 697) we conducted to gather public opinion, 83% of the people indicated that they prefer treatment in the form of probiotics or oral medication. We tested *Ec*N, a probiotic strain categorized under “generally recognized as safe” (GRAS) by the FDA [51]. It is one of the most frequently used gram-negative oral probiotic strains in research and has been studied for over a century [52]. To test if *Ec*N can be a chassis for ALDH2*1 delivery, we obtained a sample of *Ec*N (Mutaflor) from Pharma-Zentrale GmbH and attempted chemical transformation with a GFP expression plasmid (Figure 6). Although transformation efficiency was much lower compared to the common lab strain *E. coli* K12, **we still observed GFP-expressing *Ec*N colonies, indicating that transformation was successful**.

**Figure 6.**
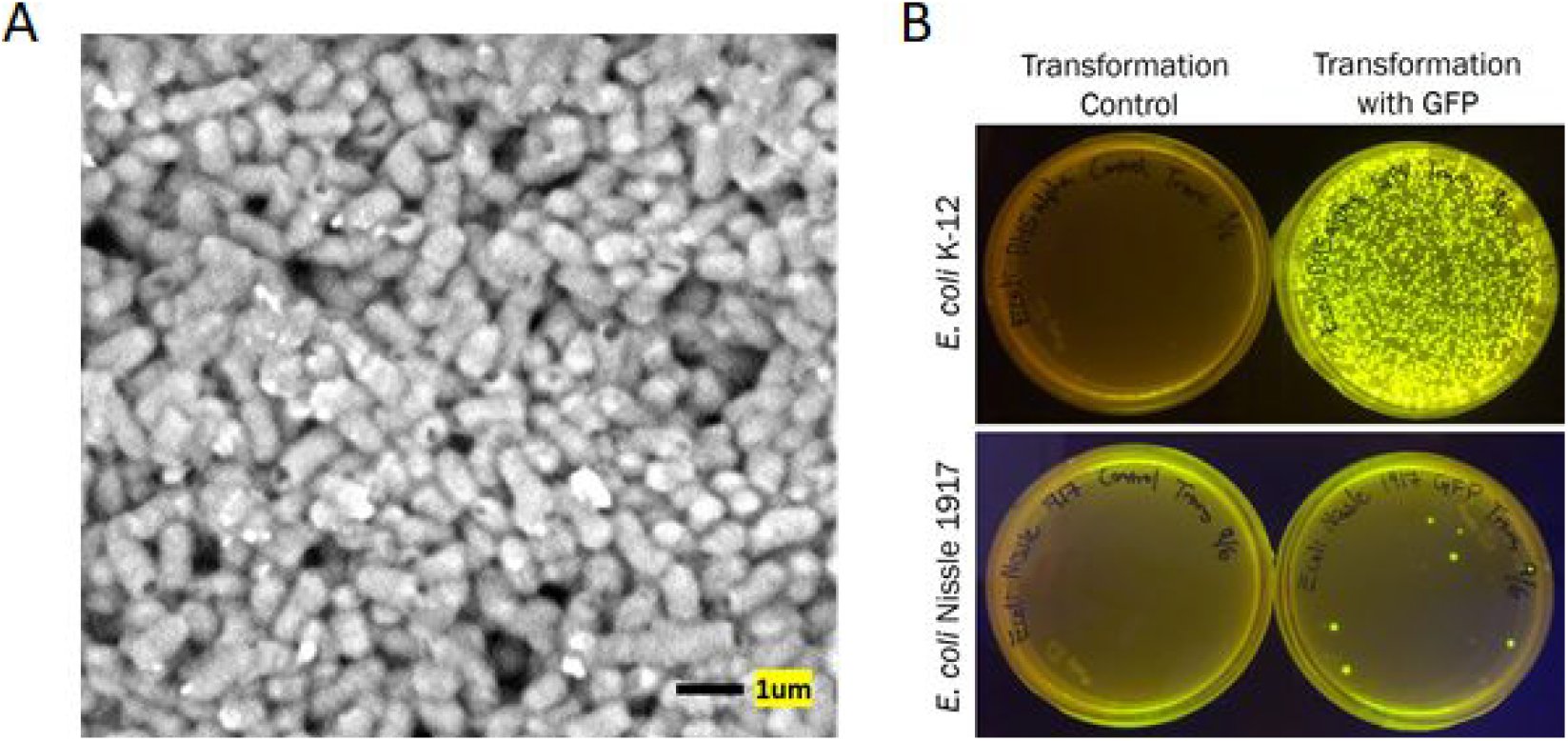
GFP was successfully transformed into *Ec*N. **(A)** *Ec*N samples were prepared for SEM by fixation with glutaraldehyde. **(B)** Both *E. coli* K-12 (**top**; positive control) and *Ec*N (**bottom**) were first treated with calcium chloride, and then chemically transformed with a GFP expression plasmid. *Ec*N colonies were glowing after transformation, showing that transformation was successful.

Next, we transformed both ALDH2*1 and ALDH2*2-expression constructs (BBa_K2539100 and BBa_K2539200), as well as a GFP positive control into *Ec*N. The transformations were successful; we observed colonies on all plates and confirmed the plasmids by PCR using the primers VF2 and VR (Figure 7). When we tested the activity of ALDH2*1-*Ec*N and ALDH2*2-*Ec*N, we observed a smaller difference in wild type and mutant ALDH2 metabolism. Since these tests were run using cell lysates, there must be other enzymes present in the cells which can also react with NAD+ or NADH; this might explain the closer levels of NADH. However, NADH levels still increased significantly faster for ALDH2*1-*Ec*N compared to ALDH2*2-*Ec*N, and did not change for the negative control (inactive ALDH2*1) (Figure 7). These results indicate that *Ec*N overexpressing ALDH2*1 can successfully metabolize acetaldehyde.

**Figure 7.**
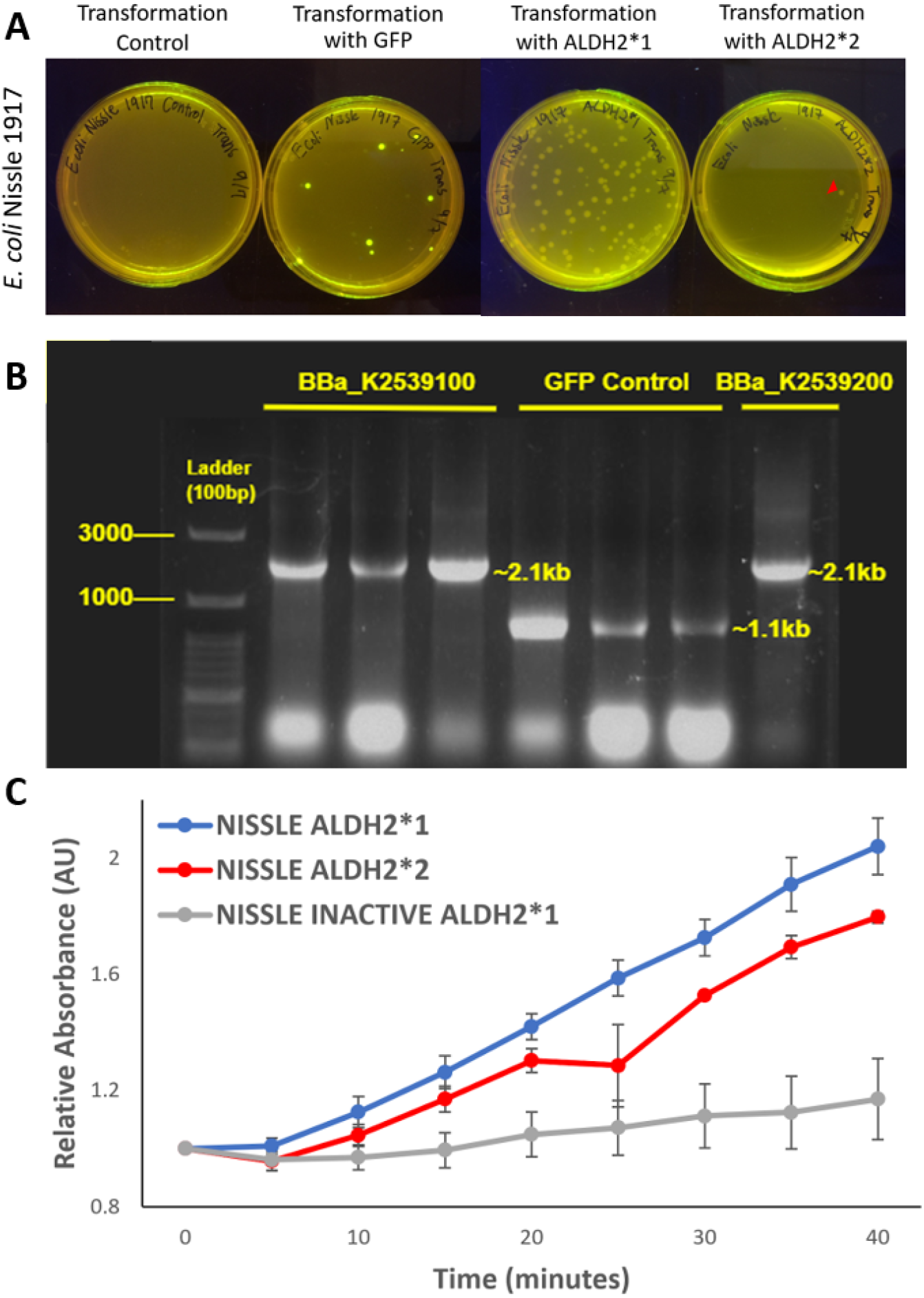
ALDH2*1-expressing *Ec*N metabolizes acetaldehyde significantly faster compared to ALDH2*2-expressing *Ec*N under simulated oral conditions. **(A)** *Ec*N was transformed with ALDH2*1 and ALDH2*2-expression plasmids (BBa_K2539100 and BBa_K2539200), and GFP as a positive control. We observed glowing green colonies for the GFP sample, as well as colonies on the ALDH2*1 and ALDH2*2 plates (red arrowhead points to the one bacterial colony of ALDH2*2). **(B)** PCR check was run using VF2 and VR primers and confirmed the transformations. We observed bands at the expected sizes: BBa_K2539100 and BBa_K2539200 are both 2.1 kb, and the GFP is 1.1 kb. **(C)** Enzymatic activity was tested using artificial saliva at 37°C. The graph shows relative activity of *Ec*N lysates containing either ALDH2*1, ALDH2*2, or inactive ALDH2*1 (boiled to denature proteins; negative control). Error bars represent standard error.

### Production and Testing of an ALDH2 Probiotics Candy

ALDH2 deficiency mainly increases the risks of developing esophageal and head and neck cancers. To directly target *salivary* acetaldehyde levels in the region, we envisioned using a candy (similar to a throat lozenge), which remains in the mouth, to steadily deliver recombinant ALDH2*1. The candy will deliver ALDH2*1 either as a purified enzyme or as ALDH2*1-*Ec*N. Here, we document the production and testing of a probiotic candy.

#### Production

The candy-making process consists of three stages: dissolving, cooling, and molding [53]. In our temperature threshold tests of GFP-*Ec*N and ALDH2*1-*Ec*N, the probiotics remained alive at temperatures lower than 60°C, so we added *Ec*N to the candy during the cooling stage (Figure 8). The final product carrying GFP-*Ec*N fluoresced, which shows that we could successfully **incorporate living probiotics carrying functional proteins into the candy**.

**Figure 8.**
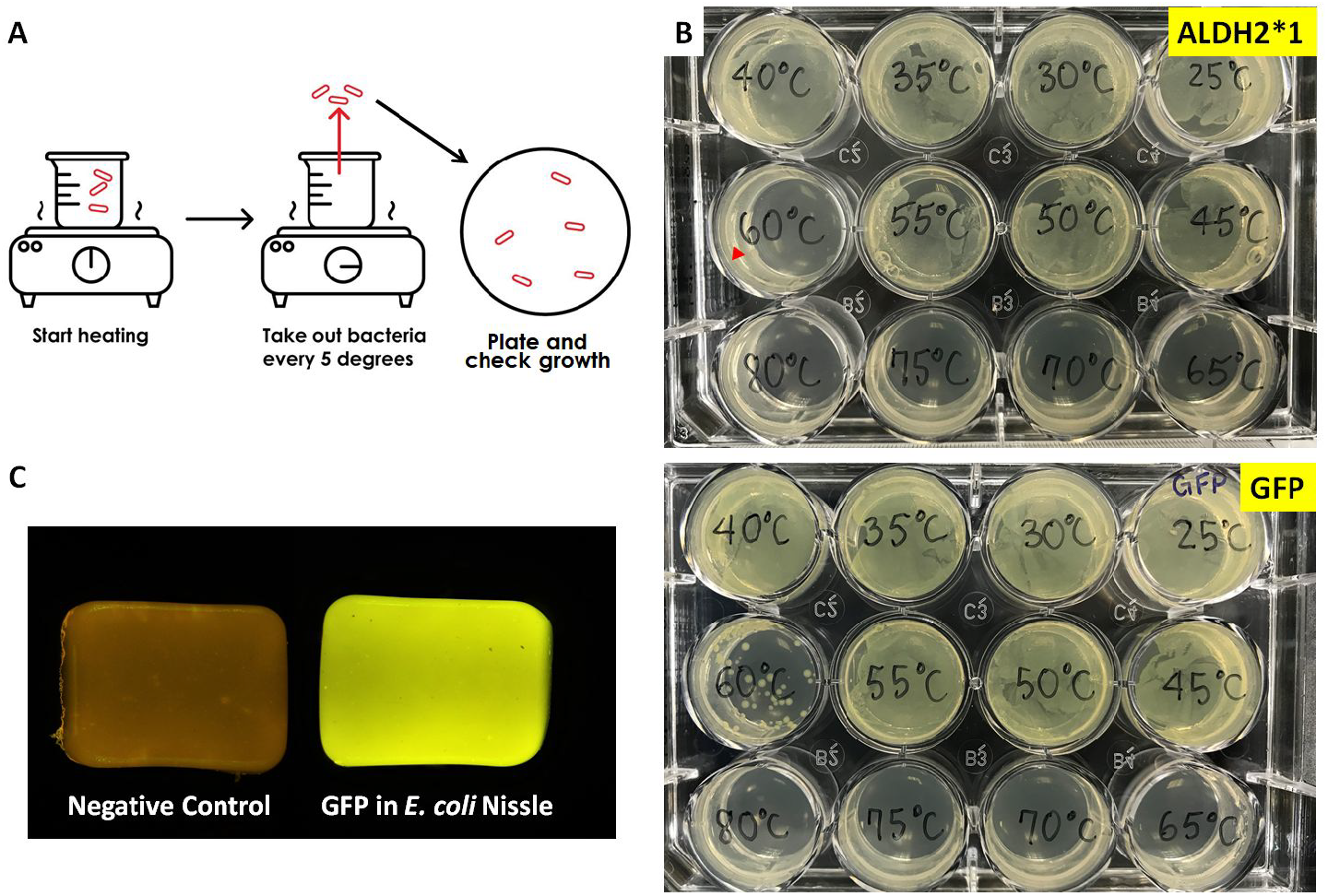
GFP was successfully expressed in an *Ec*N probiotics candy. **(A)** Testing *Ec*N growth at different temperatures; *Ec*N carrying ALDH2*1 or GFP were heated in a beaker of water and plated at 5°C increments. **(B)** Both ALDH2*1-*Ec*N (red arrowhead) and GFP-*Ec*N grow at or below 60°C. **(C)** GFP-*Ec*N candy glows under blue light, indicating that proteins are functional. The negative control does not contain any *Ec*N.

#### Testing ALDH2*1-EcN Probiotics Candy

Following the same procedure, we made ALDH2*1-*Ec*N probiotics candies and tested the candy under simulated oral conditions. As the candy dissolves, it should release ALDH2*1-*Ec*N. If this occurs too quickly, most of the ALDH2*1-*Ec*N might be swallowed right away and lose its effect in the mouth. We found that our *Ec*N candies dissolve at a rate comparable to commercial lozenges (Ricola and Nin Jiom herbal candies) in the lab. Like the lozenges, then, our *Ec*N candies should also completely dissolve after about 13 minutes in the mouth.

To test our *Ec*N candy’s ability to metabolize acetaldehyde, it was dissolved in artificial saliva, centrifuged to isolate ALDH2*1-*Ec*N, and then lysed. Enzyme activity assays were performed using cell lysates in artificial saliva at 37°C. We compared three different candies, containing either ALDH2*1-*Ec*N, ALDH2*2-*Ec*N, or no *Ec*N. Our results show that **ALDH2*1-*Ec*N released from the candy is functional and effective at metabolizing acetaldehyde** (Figure 9).

**Figure 9.**
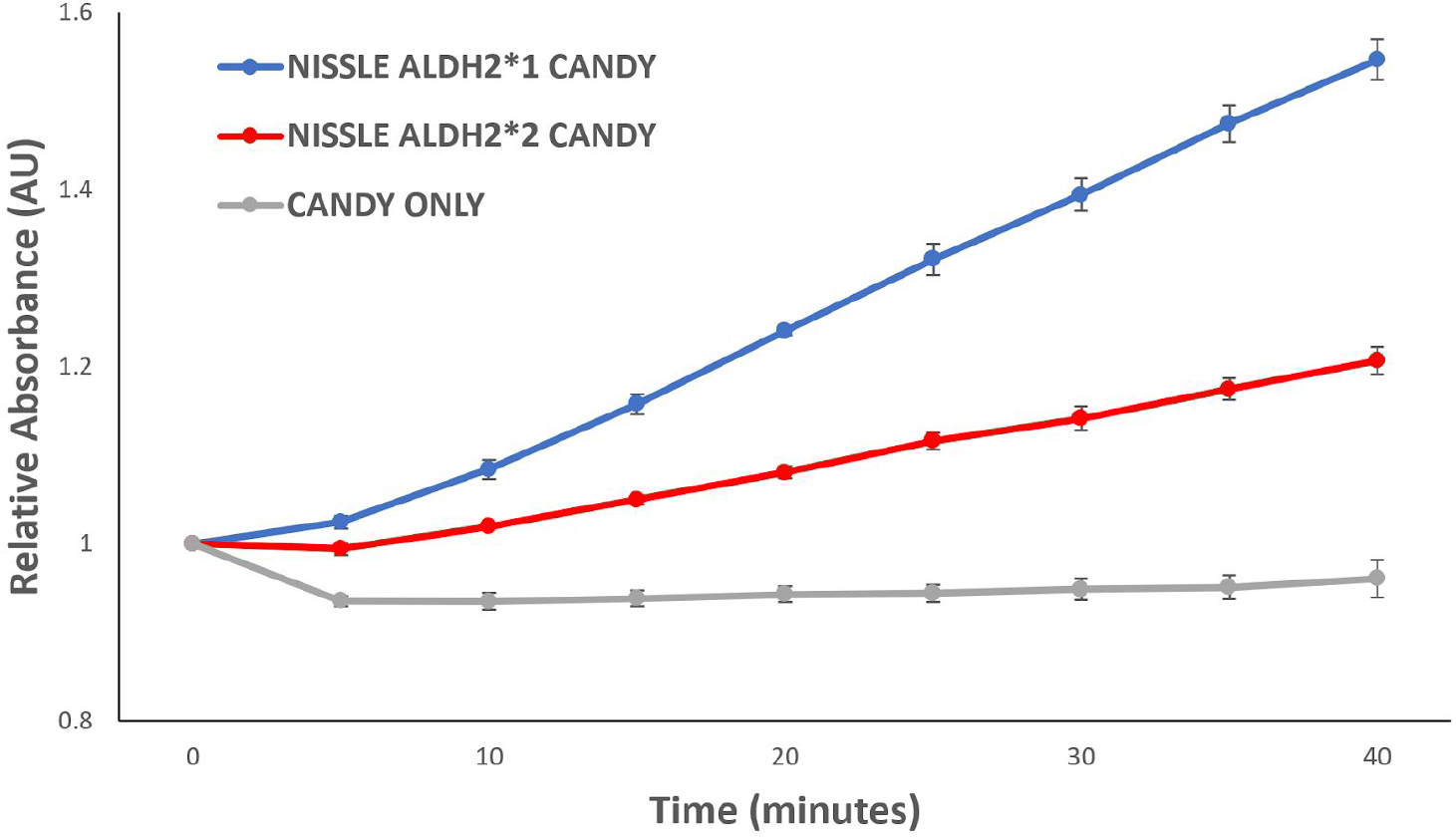
Our candy carrying ALDH2*1-*Ec*N converts acetaldehyde significantly faster than ALDH2*2-EcN. To simulate realistic conditions, the candy was first dissolved in artificial saliva at 37°C before *Ec*N cell extracts were tested. A candy that did not contain any bacteria served as the negative control. Error bars represent standard error.

### Mathematical Modeling: Supplementing ALDH2-deficient Individuals with Recombinant ALDH2*1 to Match Wild Type Enzyme Activity

For ALDH2-deficient individuals, we used mathematical modeling to calculate the amount of ALDH2*1 to consume in order to reduce salivary acetaldehyde to wild type levels. To simulate acetaldehyde concentration in the mouth over time, we first modeled the natural metabolism of alcohol as a two-step reaction (Figure 10).

**Figure 10.**
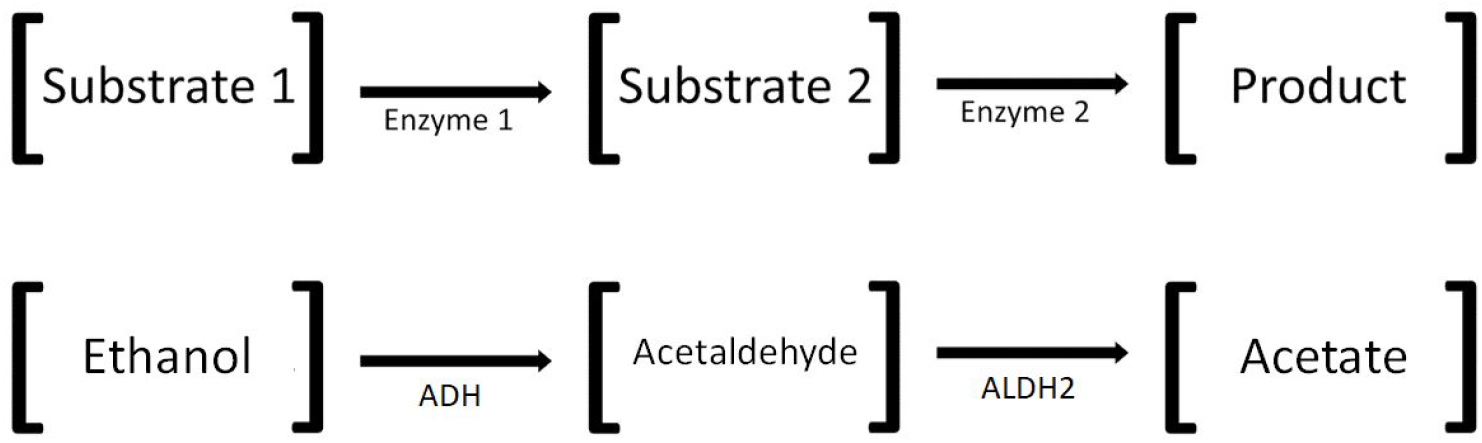
Schematic of a two-step sequential enzymatic reaction. The metabolic pathway converting ethanol to acetate occurs in two steps. To model the concentration of acetaldehyde over time, we considered the following factors: the initial concentrations of ethanol ([Substrate 1]) and acetaldehyde ([Substrate 2]), and the reaction rates of ADH (Enzyme 1) and ALDH2 (Enzyme 2).

We considered four factors to determine acetaldehyde concentrations in the mouth: initial salivary ethanol and acetaldehyde concentrations, ADH activity, and ALDH2*1 activity. We obtained literature values for salivary ethanol and acetaldehyde concentrations (Table 1) [21], as well as the reaction rate of ADH. ADH1B*2, a highly prevalent allele among East Asians (found in over 80% of Chinese, Taiwanese, Korean, and Japanese studied populations), converts ethanol to acetaldehyde at a maximum rate of 0.82 µM/sec [54], [55]. Finally, from our own experiments, we determined that at 37°C, 0.222 nM of purified ALDH2*1 enzyme converts acetaldehyde to acetate at a rate of 4.50 nM/sec (data from Figure 5).

**Table 1.** Ethanol and acetaldehyde concentrations in different alcoholic beverages and saliva. Salivary acetaldehyde concentrations were recorded after consuming one mouthful of each type of alcohol. *Salivary ethanol concentrations after 0.5 minutes were calculated as proportional to the reduction in acetaldehyde concentration after 0.5 minutes (data from [21]).

|  | Concentration in Alcoholic Beverage |  |  | Concentration in Saliva (after 0.5 min) |  |
| --- | --- | --- | --- | --- | --- |
| | Ethanol (% vol) | Ethanol (mM) | Acetaldehyde ($\mu\text{M}$) | Ethanol (mM)* | Acetaldehyde ($\mu\text{M}$) |
| **Beer** | 5 | 856 | 210 | 399 | 98 |
| **Wine** | 12 | 2053 | 1502 | 385 | 282 |
| **Spirits** | 40 | 6907 | 8824 | 349 | 446 |

With known enzyme rates and substrate concentrations, we modeled the change in acetaldehyde concentration over time using Michaelis-Menten kinetics [56] [57] according to the following equation:

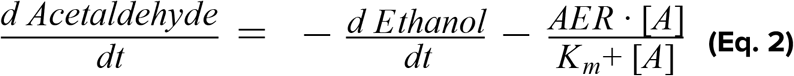

This equation relates the change in salivary acetaldehyde concentration over time to the conversion rate of ethanol to acetaldehyde (*d*Ethanol/*d*t) minus the enzymatic activity of ALDH2*1. The Michaelis-Menten constant Km for ALDH2*1 is 0.2 µM [58], and we assume the same value for each variant. The acetaldehyde elimination rate (AER) varies for different ALDH2 genotypes [3], [5]. For our model, we calculated AER for the genotypes ALDH2*1/*2 and ALDH2*2/*2 as a percentage of the wild type AER (2.1 µM/sec) [58] (Table 2). The symbol [A] represents acetaldehyde concentration.

**Table 2.** Model enzyme activities for different genotypes. The acetaldehyde elimination rate varies for different genotypes. Based on the activity levels of different genotypes compared to wild type ALDH2*1/*1 (with an AER of 2.1 µM/sec), we calculated AER for the other genotypes [3], [5], [58].

| Genotype | % Wild Type Efficiency | AER ($\mu\text{M}/\text{sec}$) |
| --- | --- | --- |
| ALDH2*1/*1 | 100 | 2.1 |
| ALDH2*1/*2 | 20 | 0.42 |
| ALDH2*2/*2 | 1 | 0.021 |

We created a Python program to numerically solve Equation 2 for the modeled two-step sequential enzymatic reaction with the LSODA method [59]. The program graphs salivary acetaldehyde concentrations over time after the consumption of various alcoholic beverages for different ALDH2 genotypes (Figure 11).

**Figure 11.**
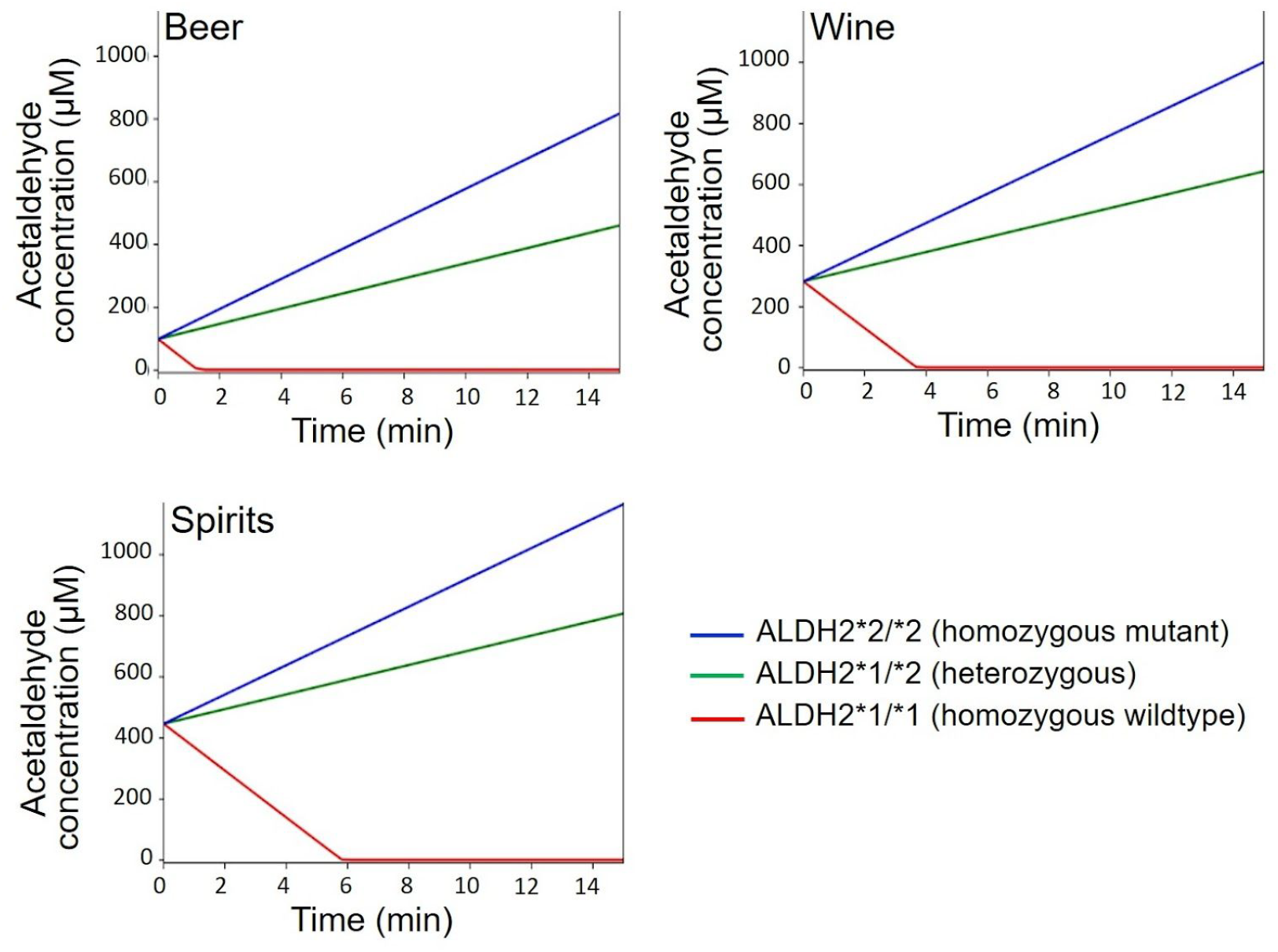
Modeled salivary acetaldehyde concentration over time after consumption of different types of alcohol (beer, wine, and spirits). The three curves in each graph represent different ALDH2 genotypes.

To determine how much additional ALDH2*1 is needed for ALDH2-deficient individuals to match normal salivary acetaldehyde levels, we modified our model to account for additional ALDH2*1 (as shown in Equation 3). The values used for wild type ALDH2*1 were determined experimentally (data from Figure 5). Manipulating the concentrations of ALDH2*1 for each genotype, we determined that a supplement of 0.103 µM ALDH2*1 would allow people with homozygous mutant ALDH2*2 to match wild type ALDH2*1 activity every minute after consuming spirits (Figure 12). The additional ALDH2*1 should reduce salivary acetaldehyde levels for ALDH2-deficient individuals, as well as the associated increased cancer risks.

**Figure 12.**
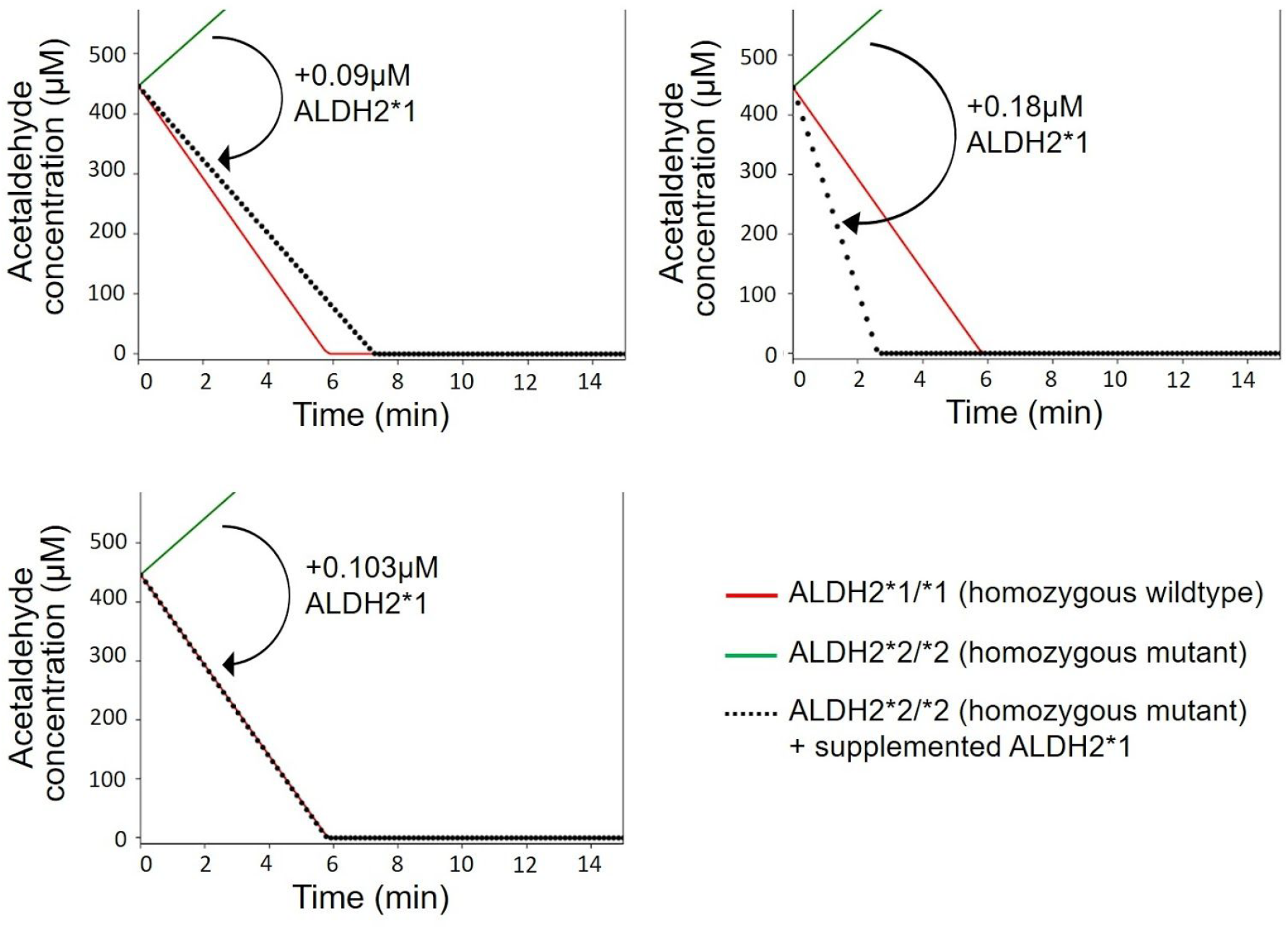
Modeled salivary acetaldehyde concentration over time after a supplement of ALDH2*1 enzymes. Here, concentrations of salivary acetaldehyde are shown after the consumption of spirits. A supplement of wild type ALDH2*1 changes the acetaldehyde levels in the mouth. To match homozygous wild type ALDH2*1/*1 activity, an additional 0.103 µM ALDH2*1 should be given to homozygous mutant ALDH2*2/*2 individuals. For other types of alcohol (not shown above), the same supplement of 0.103 µM ALDH2*1 also allows acetaldehyde levels in homozygous mutants to match those found in wild type individuals.

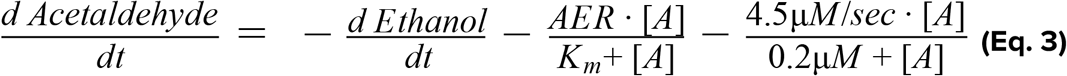

## Acknowledgments

We would like to thank Dr. Che-Hong Chen for answering our questions regarding the misconceptions about Asian glow, Dr. Stephanie Hsieh for a providing medical perspective to our project, Dr. Ying Chieh Tsai for recommending the use of *E. coli* Nissle 1917, and Dr. Rudolf von Bünau for providing us with *E. coli* Nissle 1917.

